# Capsid-E2 interactions rescue core assembly in viruses that cannot form cytoplasmic nucleocapsid cores

**DOI:** 10.1101/2021.06.28.450273

**Authors:** Julie M. Button, Suchetana Mukhopadhyay

## Abstract

Alphavirus capsid proteins (CPs) have two domains: the N-terminal domain (NTD) that interacts with the viral RNA, and the C-terminal domain (CTD) that forms CP-CP interactions and interacts with the cytoplasmic domain of the E2 spike protein (cdE2). In this study, we examine how mutations in the CP NTD affect CP CTD interactions with cdE2. We changed the length and/or charge of the NTD of Ross River virus CP and found that changing the charge of the NTD has a greater impact on core and virion assembly than changing the length of the NTD. The NTD CP insertion mutants are unable to form cytoplasmic cores during infection but they do form cores or core-like structures in virions. Our results are consistent with cdE2 having a role in core maturation during virion assembly and rescuing core formation when cytoplasmic cores are not assembled. We go on to find that the isolated cores from some mutant virions are now assembly competent in that they can be disassembled and reassembled back into cores. These results show how the two domains of CP may have distinct yet coordinated roles.

**IMPORTANCE:** Structural viral proteins have multiple roles during entry and assembly. The capsid protein (CP) of alphaviruses has one domain that interacts with the viral genome and another domain that interacts with the E2 spike protein. In this work we determine that the length and/or charge of the CP affects cytoplasmic core formation. However, defects in cytoplasmic core formation can be overcome by E2-CP interactions, thus assembling a core or core-like complex in the virion. In the absence of both cytoplasmic cores and CP-E2 interactions, CP is not even packaged in the released virions, but some infectious particles are still released presumably as RNA packaged in a glycoprotein containing membrane shell. This suggests that the virus has multiple mechanisms in place to ensure the viral genome is surrounded by a capsid core during its lifecycle.

## INTRODUCTION

Alphaviruses are enveloped viruses containing a single-stranded positive-sense RNA genome and belong to the *Togaviridae* family (1). The virion structure is comprised of three concentric shells; starting from the center of the virion these are the nucleocapsid core, a host-derived lipid membrane, and the 80 trimeric glycoprotein spikes that are embedded in the membrane (2, 3). The nucleocapsid core is composed of the viral genome surrounded by up to 240 copies of the capsid protein (CP) (2, 4). The virion’s lipid membrane is derived from the host plasma membrane where budding occurs. Each surface spike is comprised of a trimer of the E1 and E2 glycoproteins; both proteins are required for cell entry (5). E1 is a single-transmembrane protein with roughly five amino acids on the cytoplasmic side or interior of the lipid bilayer (6). The E2 protein is also a single-transmembrane protein but has a 33 amino acid cytoplasmic domain (cdE2) interior the lipid bilayer (6, 7). The cdE2 interacts with CP and this interaction is proposed to facilitate particle assembly and disassembly (2, 6, 8–13).

The CP is composed of two distinct domains; the ordered C-terminal chymotrypsin-like protease domain, and the disordered N-terminal domain (NTD) which contains a large number of positively-charged residues (14, 15). These two domains have two distinct functions. It is thought that non-specific electrostatic interactions between the basic NTD of CP and the acidic viral RNA (vRNA) facilitate assembly of nucleocapsid cores (16–19). Only a portion of the NTD is resolved in virion structures; residues 97-106 in Barmah Forest virus, residues 115-124 in Venezuelan Equine Encephalitis virus (VEEV) are modeled as an α-helix, and residues 82-112 in Eastern Equine Encephalitis virus (EEEV) are modeled as a mix of an extended coil and a short helix (7, 20, 21). The Cterminal domain (CTD) forms a lattice of hexamers and pentamers around the genome and is well-resolved in virion structures (2, 6, 7, 20, 22). An interaction between cdE2 (residues 400-402 in Sindbis virus(SINV)) and a hydrophobic pocket identified in the CTD of CP has been shown to be critical for efficient budding of infectious virus particles (8, 9, 12, 23).

Assembly and incorporation of nucleocapsid cores into budding virions is thought to occur via two proposed models (24). In the nucleocapsid (NC)-directed assembly model, NC assembly is driven by CP-RNA interactions leading to CP-CP interactions resulting in the formation of cytoplasmic cores; core assembly occurs independent of the glycoproteins. In this model, the pre-assembled cytoplasmic core imposes geometrical constraints on the budding virion, such as virion diameter, shape, and a 1:1 stoichiometry between CP and E1/E2 trimers (8, 11–13, 24–28). In the glycoprotein (GP)-directed assembly model, cytoplasmic assembly of nucleocapsid cores is not required for budding of virions. Instead, interactions between cdE2 and the hydrophobic pocket in the CTD of CP facilitates assembly and incorporation of nucleocapsid cores during budding (9, 12, 17–19, 29–31). This second model is supported by several studies with viruses containing CP mutations that alter CP-RNA and/or CP-CP interactions that prevent cytoplasmic core formation but not release of infectious virions (9, 12, 17–19, 26, 29–32).

These CP mutations include N-terminal deletions (17-19, 29, 31), N- or C-terminal substitutions (9, 12, 26, 29, 30), or C-terminal insertions (33), with differing effects on production of infectious virions from no difference to a complete loss of infectious virus release. In most cases, the mutations prevent cytoplasmic core formation as well as core formation in the virus particle (12, 19, 30, 31). However, a SINV CP mutant called TC-186 prevents cytoplasmic core formation but does form cores in the released virus particles (33). When cytoplasmic core formation is abolished in infected cells, it is thought that complexes of viral RNA and CP interact with cdE2 at the plasma membrane during budding, per the GP-directed assembly model. Both of these models rely on core assembly via RNA-CP interactions combined with either CP-CP or GP-CP interactions.

In this work, we investigated the effects of N-terminal CP insertions on cytoplasmic core and virion assembly in cells infected with the alphavirus Ross River virus (RRV). We identify mutants that are unable to assemble cytoplasmic cores in infected cells but do form core-like structures in the released virus particles, consistent with another insertion mutant (33). Furthermore, we show that core assembly in virions is dependent on CP-cdE2 interactions during the budding process. Finally, we propose that the NC- and GP-directed assembly models represent functional redundancy in alphavirus CP and CP-cdE2 interactions to facilitate production of infectious particles even when nucleocapsid assembly or incorporation into virions is compromised. However, since infectious virus release is diminished when one or both modes of nucleocapsid assembly and incorporation are abolished, it is likely that alphaviruses use both pathways to maximize release of infectious virus particles.

## RESULTS

### Capsid protein mutants to study the role of the N-terminal domain

Deletion or substitution mutations in the N-terminal domain of CP can result in the loss in the cytoplasmic cores but still produce infectious virus (9, 17–19, 29–31). Cytoplasmic cores are formed through a series of unknown intermediates following the interaction of CP with the vRNA. Since the CP NTD is responsible for interacting with the vRNA primarily through its positively-charged amino acid residues, we hypothesized that increasing the length of the CP via addition of a mixture of neutral and charged amino acids would impact cytoplasmic core formation. Addition of only neutral amino acids should not affect electrostatic interactions between the CP NTD and vRNA but could affect core formation due to increased steric hindrance inside the core.

To this end, we designed three N-terminal CP mutants to investigate the effect of additional CP length and/or negative charge on the ability of the viruses to assemble during viral infection in RRV. These mutants, D_rich_, Reverse D_rich_, and Plus25neutral, differ either in length or length and charge compared to the wild-type (WT) CP (Figure 1). In WT CP, region A consists of residues 1-82 (RRV numbering, Figure 1) and contains the previously identified coiled-coil motif (34–36). Region B consists of residues 83-107 in WT CP and has a large percentage of positively-charged residues (Figure 1). Region C consists of residues 108-121 (Figure 1) and is the highly conserved region of CP sometimes called the linker peptide (7, 19). Regions D and E are variations of region B; region D is region B with ten positively-charged amino acids mutated to negatively-charged residues, whereas region E is region B but with all charged amino acids changed to neutral amino acids (Figure 1). Therefore, Plus25neutral has 25 neutral amino acids added to the CP N-terminus and does not change the overall charge CP. However, in D_rich_ region D has been added to prior to region A resulting in a decrease in the overall charge of the CP N-terminal domain from +26 to +17. Similarly, ReverseD_rich_ changes the overall charge of the CP N-terminal domain from +26 to +17, but regions B and D are switched compared to D_rich_ CP (Figure 1). The resulting changes in molecular weight due to the longer NTD are shown in Figure 1.

**Figure 1:**
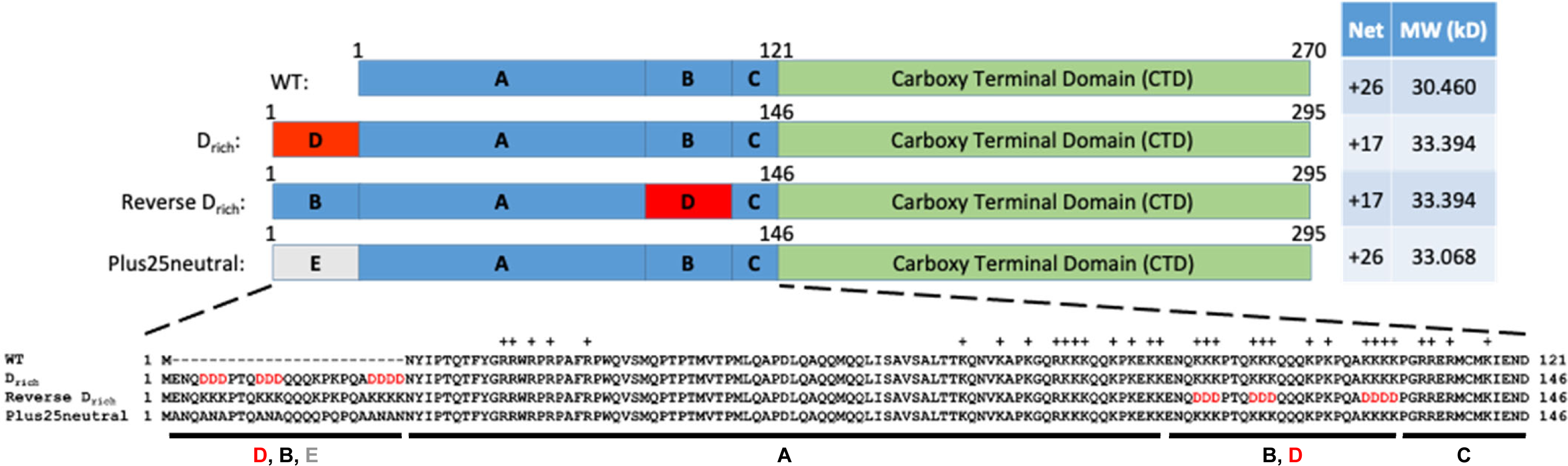
Schematics of the Ross River virus (RRV) N-terminal capsid protein (CP) mutants. The table on the right indicates the net charge of the CP N-terminus, and the molecular weight (kD) for each CP mutant. The sequences for the N-terminus of each CP mutant are indicated below the schematic with residues mutated to aspartate in red.

### CP is expressed in all of the CP mutants but cytoplasmic core formation is abolished

First, we determined protein expression and cytoplasmic core formation in our mutants compared to WT RRV. Cells were infected at an MOI=1 for 24 hours, and then cells were lysed, run on an SDS-PAGE gel and probed by western blot for CP and E1/E2 expression. We found that all of the viruses express the structural proteins although there may be slightly less expression for the D_rich_ CP than the other capsid proteins (Figure 2A). Next, we wanted to know if the increased length and/or charge would impact cytoplasmic core assembly during a viral infection. Cells were infected at an MOI=1 for 24 hours and then lysed with a non-denaturing detergent. The lysates were run over a sucrose gradient, fractions collected and run on an SDS-PAGE gel, and the viral proteins detected using trichloroethanol (37). We found that only WT virus forms cytoplasmic cores. These cores migrate to the bottom half of the gradient (Figure 2B, red box) as has been previously shown in the literature (13, 29, 38). However, we were unable to detect cores in the cells infected with CP mutant viruses (Figure 2B). These results show that N-terminal CP mutations that increase CP length, can impact cytoplasmic core formation.

**Figure 2:**
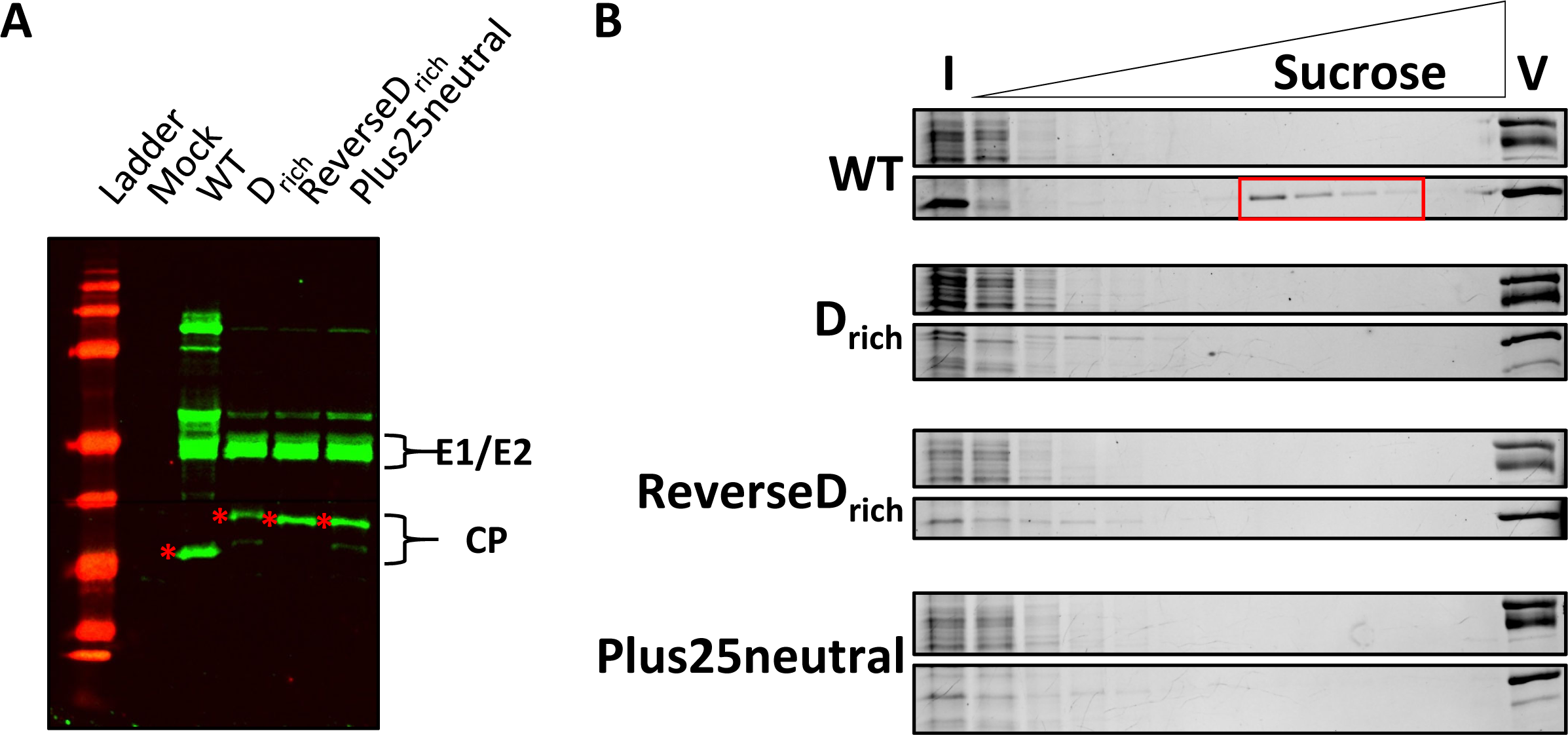
The capsid proteins are expressed in the mutant strains, but only WT virus is able to assemble cytoplasmic cores during an infection. (A) Cell lysate western for BHK cell lysates when cells were infected at an MOI=1 for 24 hours. Viral proteins were detected using α-CP and α-E1/E2 primary antibodies followed by goat antirabbit AlexaFluor^TM^750 secondary antibody. The asterisks indicate the expected capsid protein band for each virus. (B) Cytoplasmic cores were isolated from BHK cells infected at an MOI=0.1 for 24 hours as described in the Materials and Methods section. The cores were fractionated over a 10-40% w/v sucrose gradient, and equal volumes of each fraction were loaded on 12% SDS-PAGE gels containing TCE. Gels were imaged using the stain-free method; the gel images are representative of at least three biological replicates. I= input; V= virus control.

### The CP mutations do not affect viral titers but do affect the size of released virus particles

Previous work has shown cytoplasmic cores are not necessary for assembly of infectious virions. We wanted to examine if our mutant viruses released infectious virions (Figure 3). When BHK cells were infected at an MOI=1 for 24 hours, we found that the titers of viral supernatants for all viruses were within a half a log of each other (Figure 3A). This was surprising to us but not unprecedented. It has been shown that CP mutants that are unable to form cytoplasmic cores are still able to produce infectious virus, although titers are reduced compared to WT virus (9, 12, 17–19, 29–31). It has also been shown that glycoproteins can package vRNA in the absence of CP, forming infectious microvesicles (iMVs) that have a several log reduction in titer compared to WT virus (32). To determine if CP was incorporated into the infectious particles, media from infected cells was purified through a sucrose cushion, resuspended, and analyzed on an SDS-PAGE gel. We found that all of the mutant CPs are being packaged into the viral particles along with E1 and E2, suggesting that the CP and vRNA are still enveloped by the glycoproteins and incorporated into the budding virus particles (Figure 3B). Interestingly, for the D_rich_ and Plus25neutral viruses, there appears to be two CP bands, one at the expected molecular weight, as well as a lower band at approximately the molecular weight of WT CP (Figure 3B); this was also observed in infected cell lysates (Figure 2A).

**Figure 3:**
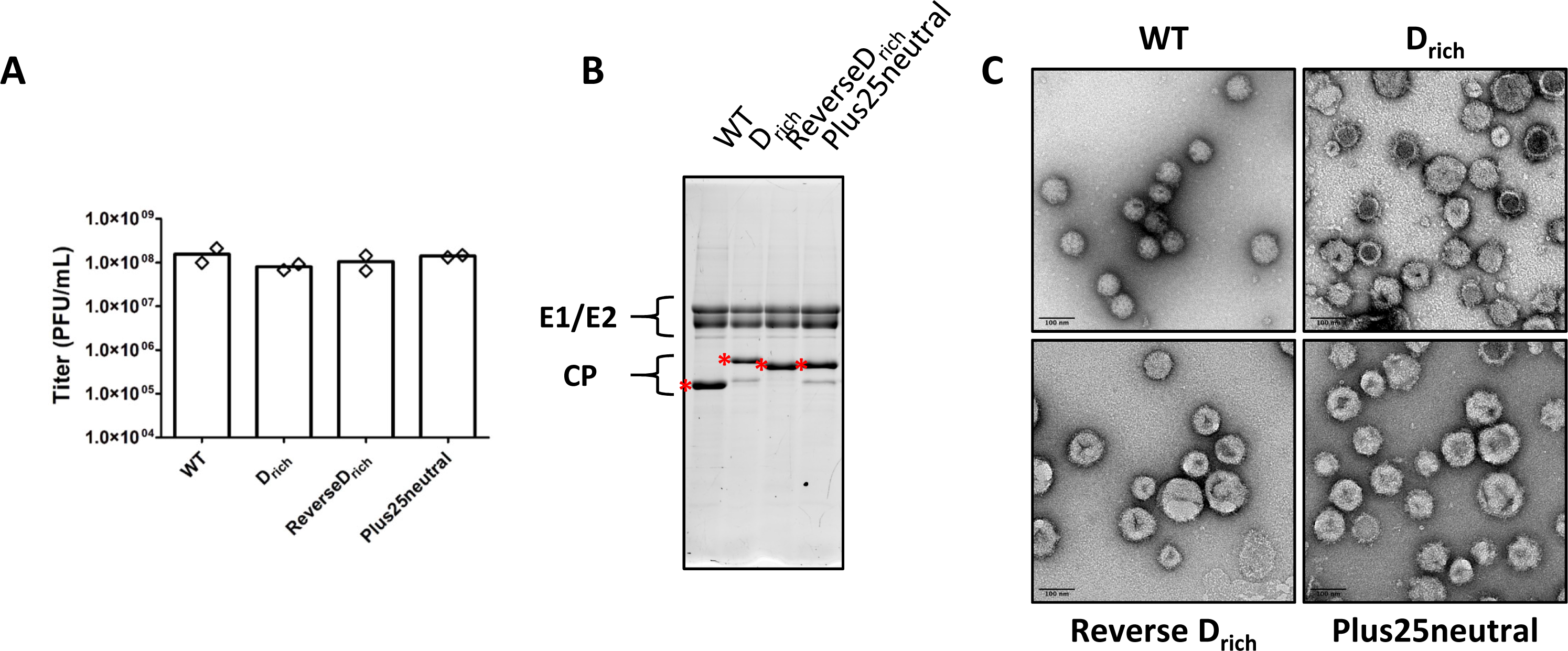
The N-terminal CP mutations do not affect viral titers but result in altered particle morphologies. (A) Viral titers as determined by plaque assay at 24hpi when infecting BHK cells at an MOI=1. The results are shown as the average (bar) of two biological replicates (diamonds). (B) Virus particles were purified by pelleting through a 27% w/v sucrose cushion and then detected on a 12% SDS-PAGE gel using TCE. The asterisks indicate the expected capsid protein band for each virus. (C) Virus particles purified by pelleting through a sucrose cushion were imaged via negative-stain TEM. Scale bar= 100nm.

We wanted to examine if the size and morphology of the released mutant virus particles was altered compared to WT RRV. Recent computational work has indicated that the benefit of core formation for alphaviruses is that it results in particles of uniform size and morphology (24), so we hypothesized the mutant virus particles would be less uniform in size and shape. When we look at the particles by negative-stain TEM we found that the mutant viruses produced particles of varying sizes (Figure 3C). Consistent with WT being able to form cytoplasmic cores, TEM images show that WT virus produces uniform spherical particles of approximately 70nm diameter (Figure 3C, top left panel). In contrast, the mutant virus particles are approximately 70-150nm in size (Figure 3C, top right and bottom panels). These particles also appear to have defects including dark internal staining and stained “creases” suggesting that they may have collapsed during the staining process (Figure 3C, top right and bottom panels). This is consistent with a lack of or reduced number of cytoplasmic cores present during the assembly process. We know however, that RNA and CP are present in these purified particles from titer and SDS-PAGE data (Figure 3A and 3B).

### All virus particles migrate similarly on a gradient despite differences in size

The mutant virus particles were found to have a wide variety in size, so we anticipated that the mutant virus particles would migrate differently than WT particles on a sucrose gradient since migration in a sucrose gradient is based on both density and size. Instead, what we found is that all of the virus particles, mutant and WT, migrated similarly (Figure 4, top). Although particles migrated to the same area of the gradient, the proportion of particles in each fraction appears to change slightly between viruses (Figure 4, top). For the WT and D_rich_ viruses, the fraction with the most total particles (based on protein band intensity) matches the fraction with the most infectious particles (Figure 4, bottom). However, for Reverse D_rich_ and Plus25neutral, the number of total particles peaks higher in the gradient than infectious particles, indicating that there is a higher fraction of noninfectious particles higher up in the gradient (Figure 4, bottom). It is unclear whether these slight differences in migration of infectious virus particles as determined by plaque assay compared to total virus particles as determined by protein signal has a biological effect during the viral lifecycle.

**Figure 4:**
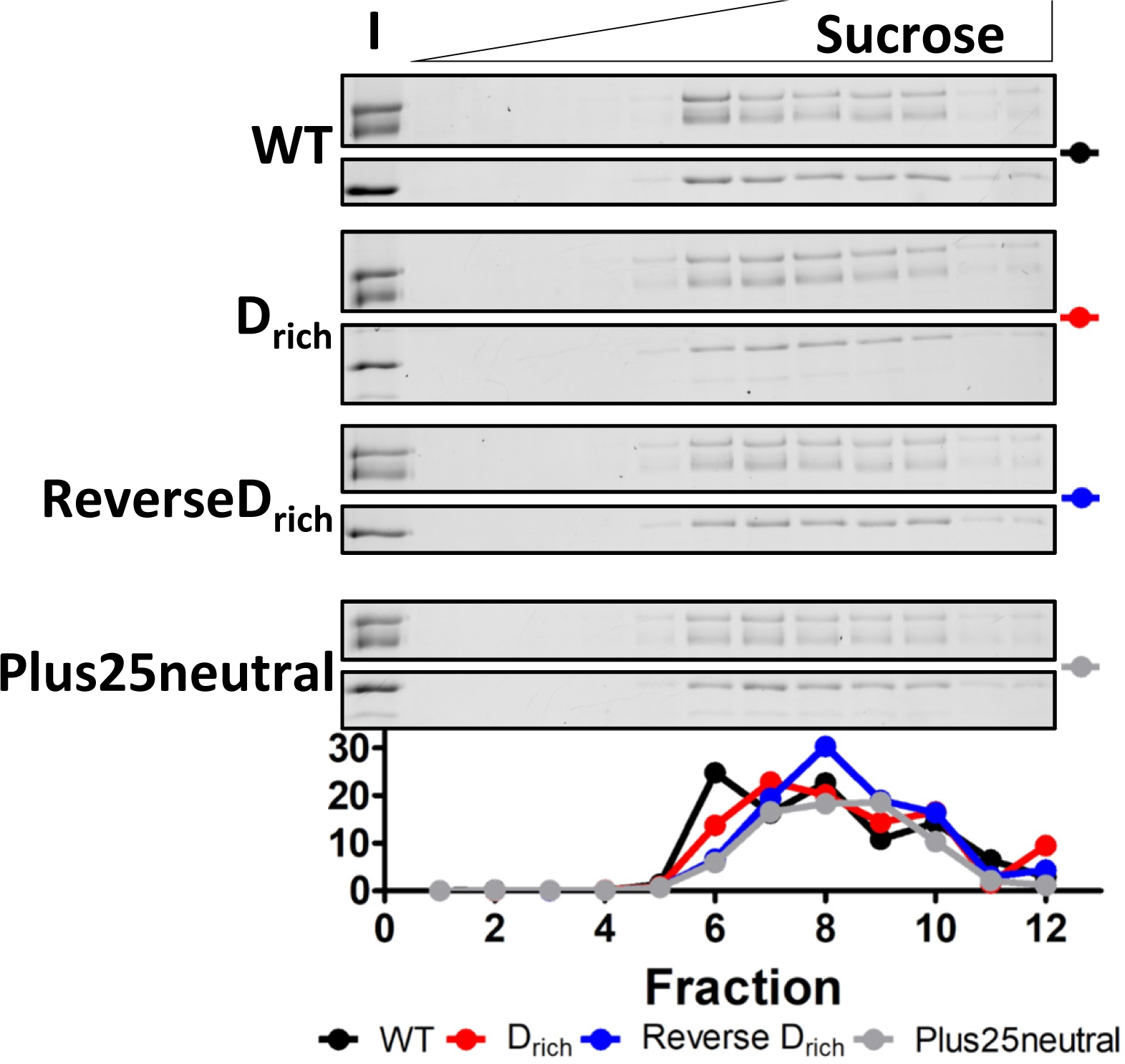
Differences in particle morphology do not appear to affect migration of virus particles through a sucrose gradient. Virus particles were pelleted through a 27% w/v sucrose cushion followed by fractionation over a 20-50% w/v sucrose gradient. Equal volumes of each fraction were loaded on 12% SDS-PAGE gels and imaged using TCE. I=input. Below the gels, titers given as a percentage of total titer for each gradient are provided. Titers were determined on BHK cells as described in the Materials and Methods section. The gel images are representative of two biological replicates, while the bottom graph is the average of two biological replicates.

### Cores are isolated from mutant virus particles

We have shown that cytoplasmic core formation for the N-terminal CP mutants is hindered in infected cells (Figure 2B), but that the CP and RNA are still packaged in the released virus particles (Figures 3A and 3B). The released particles have varied morphologies suggesting no consistent core size (Figure 3C) but the density of these particles is similar (Figures 4). One explanation is that the mutant capsid proteins were able to form cores or core-like structures after being packaged in the virus particles. To test this in our CP mutants, we purified virus particles, stripped the glycoproteins from each virus, and purified the remaining cores on the same sucrose gradient used to isolate cytoplasmic cores (Figure 2B). The cores from WT virus particles migrated in the same region of the gradient as the cytoplasmic cores (Figures 5 and 2B) and no glycoproteins were present demonstrating that the glycoproteins had been removed. As shown in Figure 2B, none of the CP mutants formed cytoplasmic cores, but the CP signal for these mutants is present throughout much of the gradient for the cores isolated from virus particles (Figure 5). All three mutants demonstrated different CP migration in the gradient for cores from particles versus cytoplasmic cores (Figures 5 and 2B). Based on migration, it appears that the Plus25neutral mutant forms cores most similar to WT cores, suggesting that increasing the length of the CP has less of an affect than also changing the overall charge of the CP.

**Figure 5:**
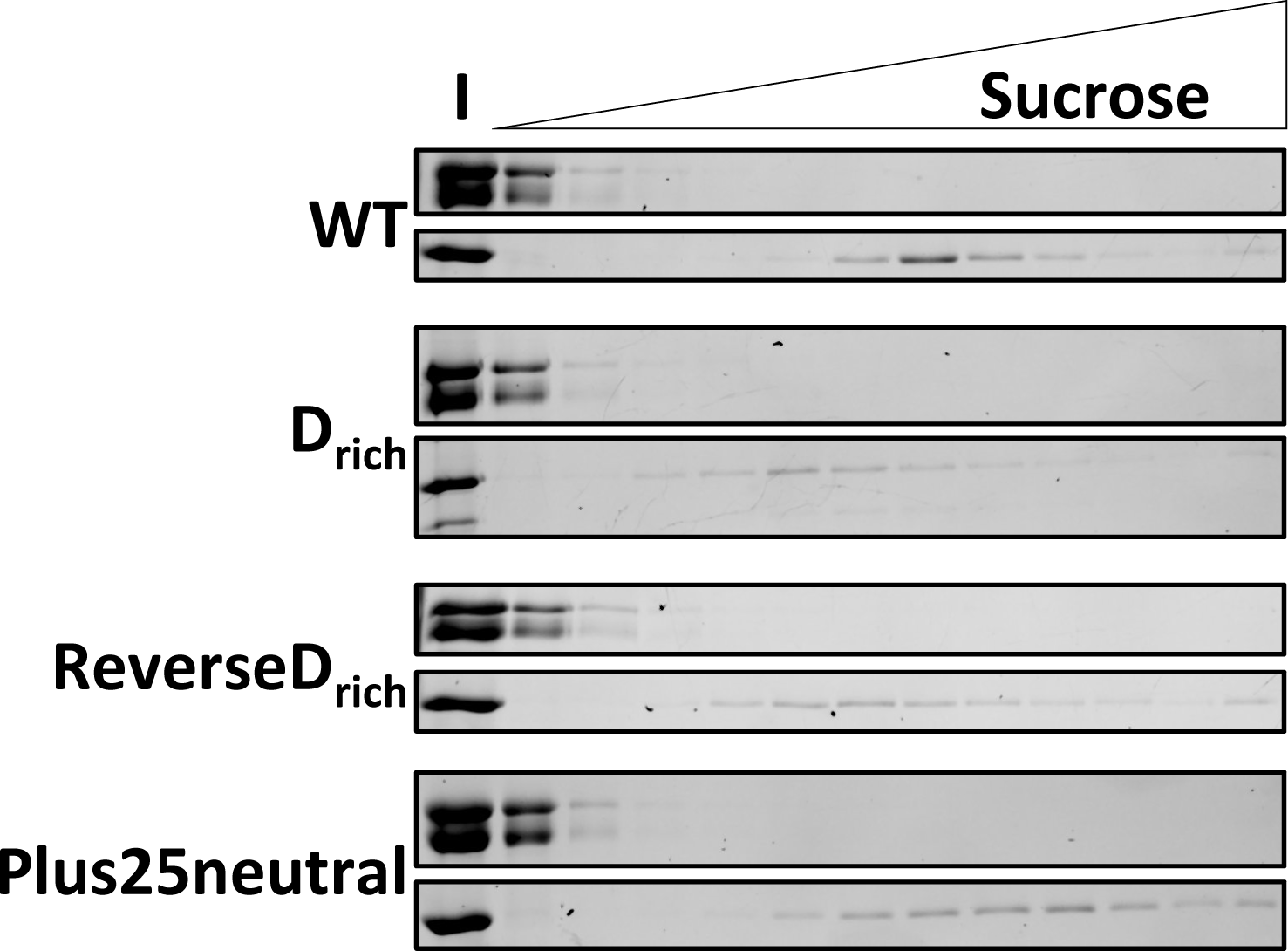
Cores isolated from virus particles for the mutant viruses migrate differently on a sucrose gradient compared to cytoplasmic cores. Cores were isolated from purified virus particles as described in the Materials and Methods section, then separated over a 10-40% w/v sucrose gradient. Equal volumes of each fraction were loaded on 12% SDS-PAGE gels and imaged using TCE. The gel images are representative of at least three biological replicates. I=input.

Overall, these results suggest that a conformational change occurs during or after virus budding. These changes result in formation of core or core-like structures that remain stable even after the glycoprotein layer is removed. Next, we had two questions: are the cores packaged in the mutant viruses similar to the cores in WT particles, and second, is core formation in virions dependent on the cdE2 interaction with CP? As the CP-cdE2 interaction is known to be a crucial part of budding, we hypothesized that this interaction may drive core formation by modulating RNA-CP and/or CP-CP interactions as budding occurs or once budding is completed.

### WT and Plus25neutral virion cores can assemble *in vitro*

*In vitro* core-like particle studies have shown us that WT cores will disassemble in the presence of high salt and then reassemble when ionic strength is reduced (4, 39, 40). Using the *in vitro* work as foundation, we developed a core disassembly and reassembly assay to characterize the cores from the mutant CP virions. Here, cores from virions were isolated (as in Figure 5), purified, treated in high salt to disassemble and then the salt concentration was reduced to promote re-assembly of cores. This assay would address if cores inside virions were similar and how length and/or charge of the CP might affect core assembly.

To isolate the cores/core-like structures from virions, virus particles were purified through a sucrose cushion, the glycoproteins removed, and the isolated cores further purified through a second sucrose cushion. Each step of core isolation (input, top of cushion, sucrose cushion, and the core pellet) was run on an SDS-PAGE gel to ensure the cores were in fact separated from the glycoproteins and could be pelleted through the sucrose cushion (Figure 6A). We found that the cores from the mutant viruses could be purified although the amounts recovered varied (Figure 6A, red asterisks). The isolated cores were imaged by negative-stain TEM (Figure 6B). The WT cores appear to be spherical and the most uniform in size and structure (Figure 6B). The cores for the mutant capsid proteins also appear to be relatively spherical but have more variation in size (Figure 6B).

**Figure 6:**
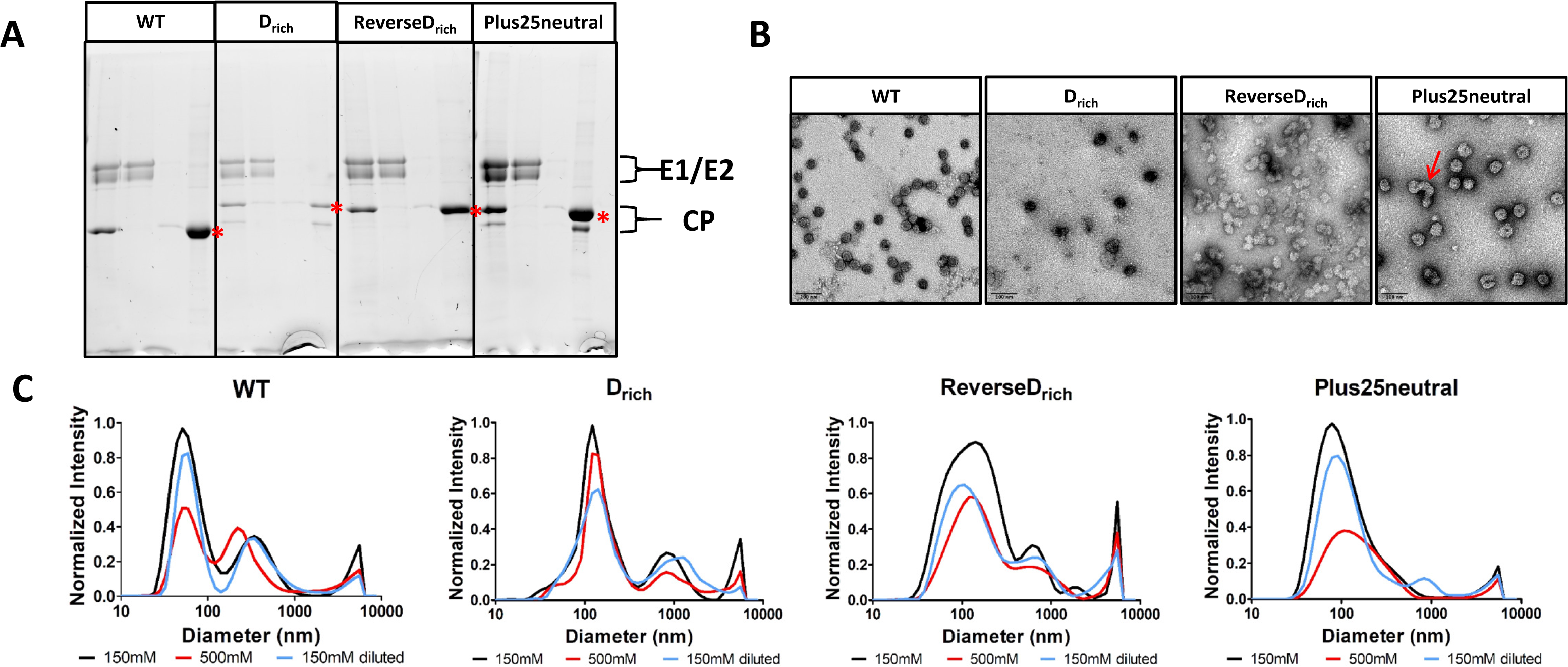
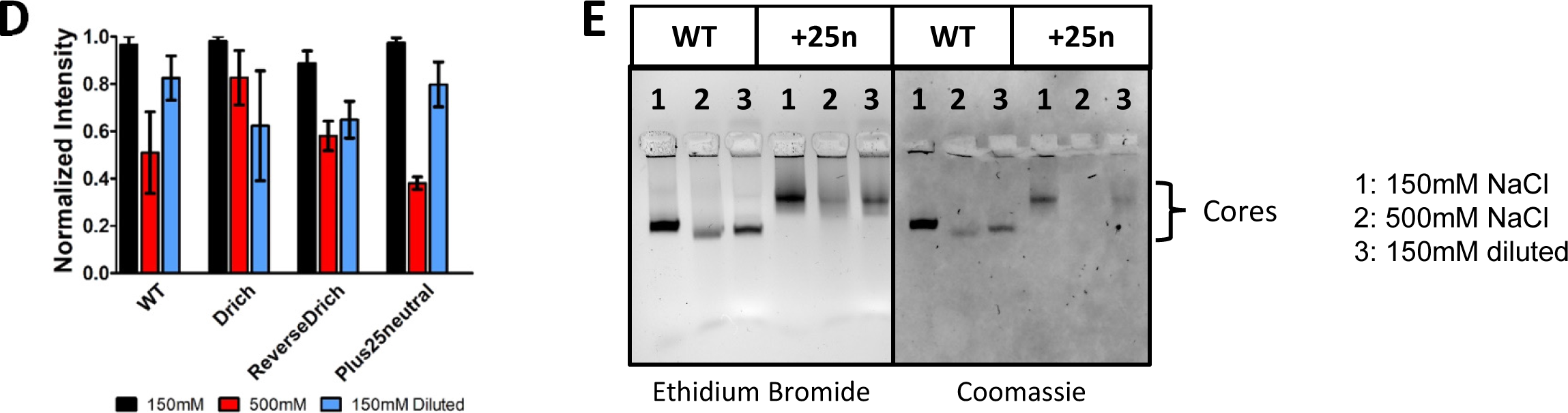
Only one mutant CP is able to reassemble cores in the absence of E2 in an *in vitro* disassembly and reassembly assay. (A) SDS-PAGE gels demonstrating the ability to pellet cores. For each virus, the lanes are in the following order (1) input, (2) top of cushion, (3) cushion, and (4) pellet. The samples were run on a 12% SDSPAGE gel and detected using TCE; the gel is representative of three biological replicates. (B) Negative-stain TEM images of the core pellets. Scale bar= 100nm. (C) Dynamic light scattering (DLS) data for the core disassembly and reassembly assay. The data is the average of two to three biological replicates and has been normalized based on counts, intensity and dilution factor as described in the Materials and Methods section. (D) Averaged DLS data from the highest peak in (C) for each sample and condition tested. (E) Agarose gel-shift assay of the WT and Plus25neutral samples in (C) demonstrating reassembly of cores following dilution to 150mM salt.

Isolated cores were disassembled using high salt (500mM NaCl) and then reassembled by diluting the salt concentration back down to 150mM. Disassembly and assembly were assayed using dynamic light scattering (DLS) and native agarose gel shift assay. What we found is that WT cores are able to disassemble and reassemble as expected in the assay based on DLS and gel shift data (Figures 6C-6E). This is based off of the decrease in amount of cores (peak at ∼50nm) from 150mM (black) to 500mM (red) NaCl, and then an increase in the amount of cores from 500mM to the diluted 150mM NaCl sample (blue) when accounting for dilution. Larger species are visible by DLS (Figure 6C) but are not clearly visible by TEM (Figure 6B), suggesting that the larger peaks are likely aggregates of cores that are only found in solution. Surprisingly, the Plus25neutral mutant was also able to disassemble and reassemble into assemblies, although the structures are larger than WT cores (Figures 6C-6E). The ReverseD_rich_ mutant was also able to disassemble to an extent and slightly reassemble, but again, the structures are larger than WT cores (Figures 6C and 6D). In contrast to these cores, the D_rich_ CP mutant actually disassembles more when diluted to lower salt concentrations (Figures 6C and 6D). Taken together, Plus25neutral is similar to WT CP cores suggesting that adding neutral amino acids to the CP doesn’t severely affect core formation in virions albeit these virions and isolated cores are larger as evident from TEM (Figure 3C and Figure 6B). In contrast, the addition of negatively-charged residues does affect core formation and even though these proteins are in virions, they are different than WT CP cores.

### The interaction of cdE2 with CP is necessary for packaging of mutant CPs into virus particles

The CP-cdE2 interaction is known to be a crucial part of particle budding (8, 9, 12, 23). We hypothesized that the interaction of CP with the E2 glycoprotein causes conformational changes resulting in formation of cores in the virus particle when core formation is hindered in the cytoplasm. If this interaction is abrogated, then viruses that cannot form cytoplasmic cores will still not be able to form cores in virus particles. To test this, we mutated the YAL residues in cdE2 to AAA. Previous work has shown that this mutation results in a significant loss in the release of infectious virus, with total virus output affected to a lesser extent (8). Our mutant CP and WT CP viruses with the cdE2 AAA mutation all had a reduction in titers compared to when cdE2 was YAL. We see a decrease in titer ranging from 4-5 logs (Figure 7A). The CP mutant viruses appeared to be more negatively affected by the addition of the E2 mutation than WT virus, which correlates with their inability to form cytoplasmic cores.

**Figure 7:**
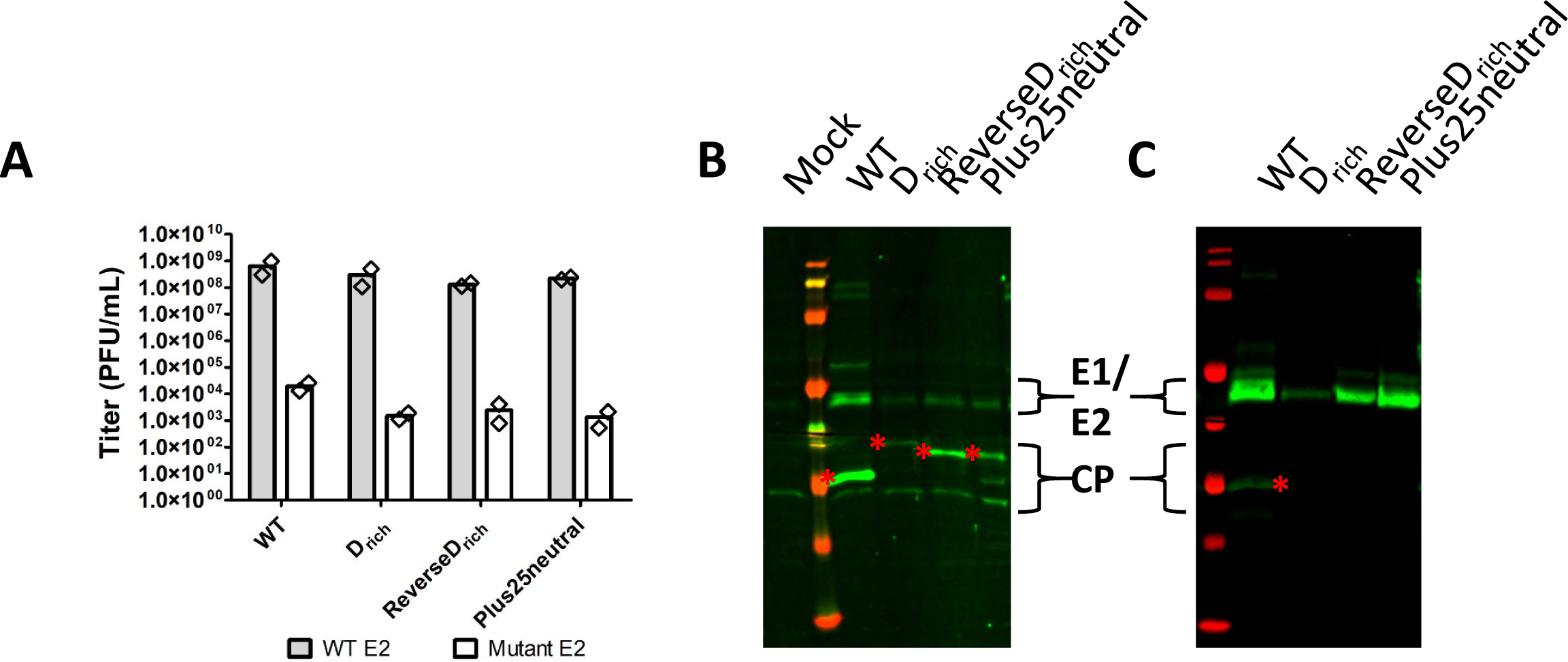
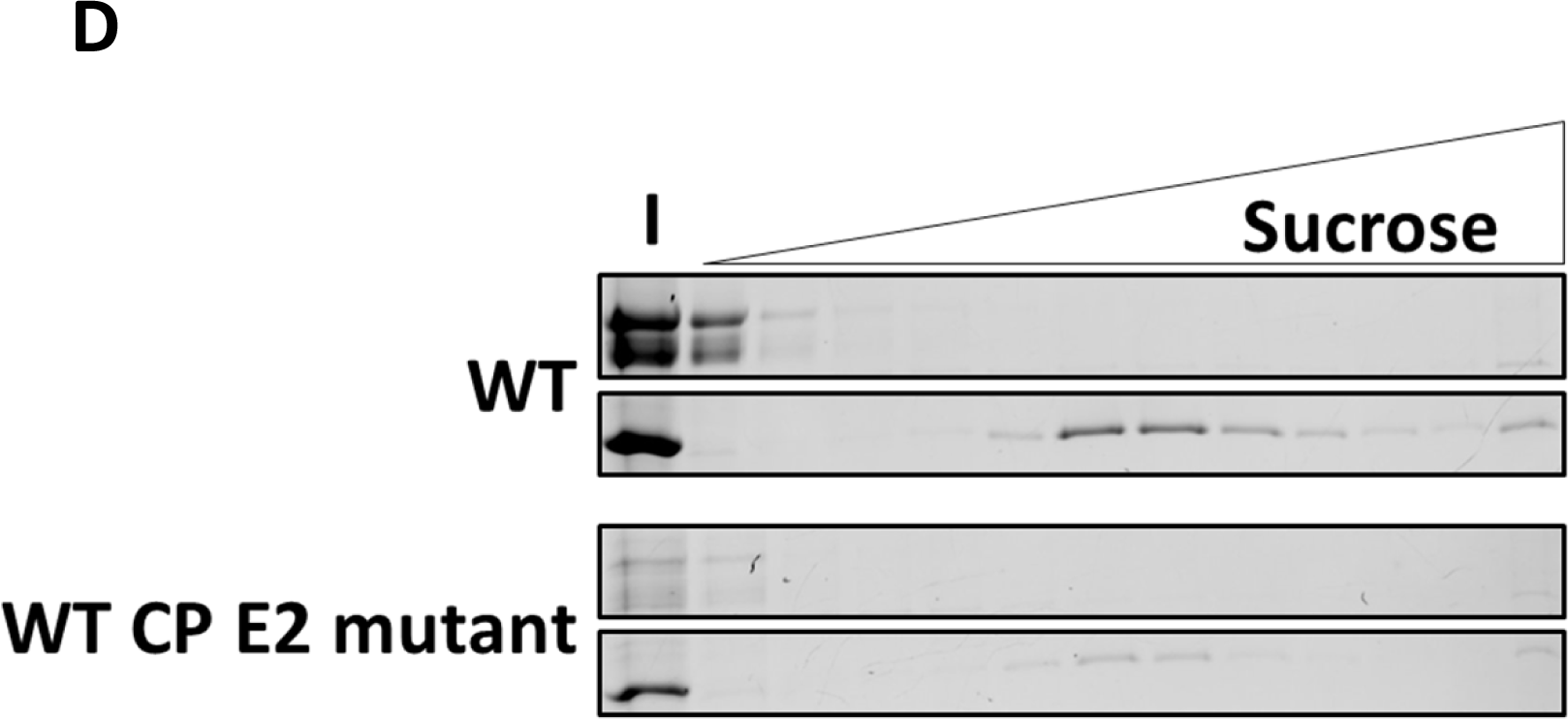
When cdE2-CP interactions are diminished in the presence of the N-terminal CP mutations, packaging of CP decreases for the CP mutants. (A) Viral titers as determined by plaque assay at 48hpi when infecting BHK cells at an MOI=0.0001. The results are shown as the average (bar) of two biological replicates (diamonds). (B) Western blot for BHK cells lysates infected at an MOI=0.0001 for 48 hours. Viral proteins were detected using α-CP and α-E1/E2 primary antibodies followed by goat antirabbit AlexaFluor^TM^750 secondary antibody. The asterisks indicate the expected capsid protein band for each virus. (C) Western blot of virus particles purified through a sucrose cushion. Viral proteins were detected as in (B). (D) Core isolations from virus particles for WT virus and the virus with WT CP but a mutated E2 protein. Cores were isolated from purified virus particles as described in the Materials and Methods section and then separated over a 10-40% w/v sucrose gradient. Equal volumes of each fraction were run on a 12% SDS-PAGE gel and proteins were detected using TCE. I=input.

When we look at the cell lysates for the E2 mutant viruses, we find that the CP and glycoproteins are expressed to varying extents (Figure 7B). WT CP has the highest amount of expression which correlates with release of more infectious virus particles.

Although there is less expression for the other CP mutants, all capsid proteins are expressed in cells infected with the E2 mutant viruses. However, in released virus particles there were differences in packaging of the capsid proteins. We found that even though all of the different viruses express CP, only the WT virus packages CP in the released virus particles (Figure 7C). Since the virus with a WT CP but a mutant E2 protein still packages CP, we tested whether the packaged CP was in a core structure. To test this, the virus was purified through a sucrose cushion, the glycoproteins stripped, and the cores isolated over a sucrose gradient (Figure 7D). Using the WT virus containing both a WT CP and WT E2 as a control, we found that the WT virus with a mutant E2 still packages cores in the virus particles (Figure 7D). These results are consistent with our hypothesis: only capsid proteins that form cytoplasmic cores will still package cores in the virus particles when the CP-cdE2 interaction is abrogated.

## DISCUSSION

### Nucleocapsid core formation is not necessary for release of infectious virions

In the traditional nucleocapsid-directed assembly pathway, CP interacts with RNA in the cytoplasm to form cytoplasmic cores which then interact with cdE2 at the membrane resulting in the release of infectious virions of uniform size and morphology (Figure 8A). This is seen with the virus that contains both WT CP and WT E2 in this study (Figures 2 and 3). In previous studies where a mutant CP results in the inability to form cytoplasmic cores, glycoprotein-directed assembly has been proposed to explain how CP and RNA is still packaged into the virus particles (Figure 8B). When our CP insertion mutants were examined in the context of viral infection, we found that core formation is abrogated but infectious virus is still released, consistent with this model (Figures 2 and 3). This result occurs whether the increased CP size is from the addition of neutral amino acids or a mixture of neutral and charged amino acids. This result along with previous observations with CP deletion mutants suggest that alphaviruses have optimized the CP length and charge to form cytoplasmic cores. Based on the inability of deletion mutants in the C-terminal end of the CP N-terminus in Semliki Forest virus (SFV) to form cytoplasmic cores, it has been proposed that this region of CP (region C in Figure 1) is necessary for cytoplasmic core formation (19, 29). Since our CP mutants contain this region, our results would suggest that this region is necessary but likely not sufficient for the formation of cytoplasmic cores. It is possible that in the mutants described in this paper, the CP orientation is altered so that this proposed core forming region is no longer able to interact and form cores. Additionally, the mutants containing additional negatively-charged residues could have altered CP-RNA interactions since it is thought that positively-charged residues in the CP N-terminus mediate CP-RNA interactions. These mutations could affect the ability of capsid proteins to form the necessary interactions for core formation.

**Figure 8:**
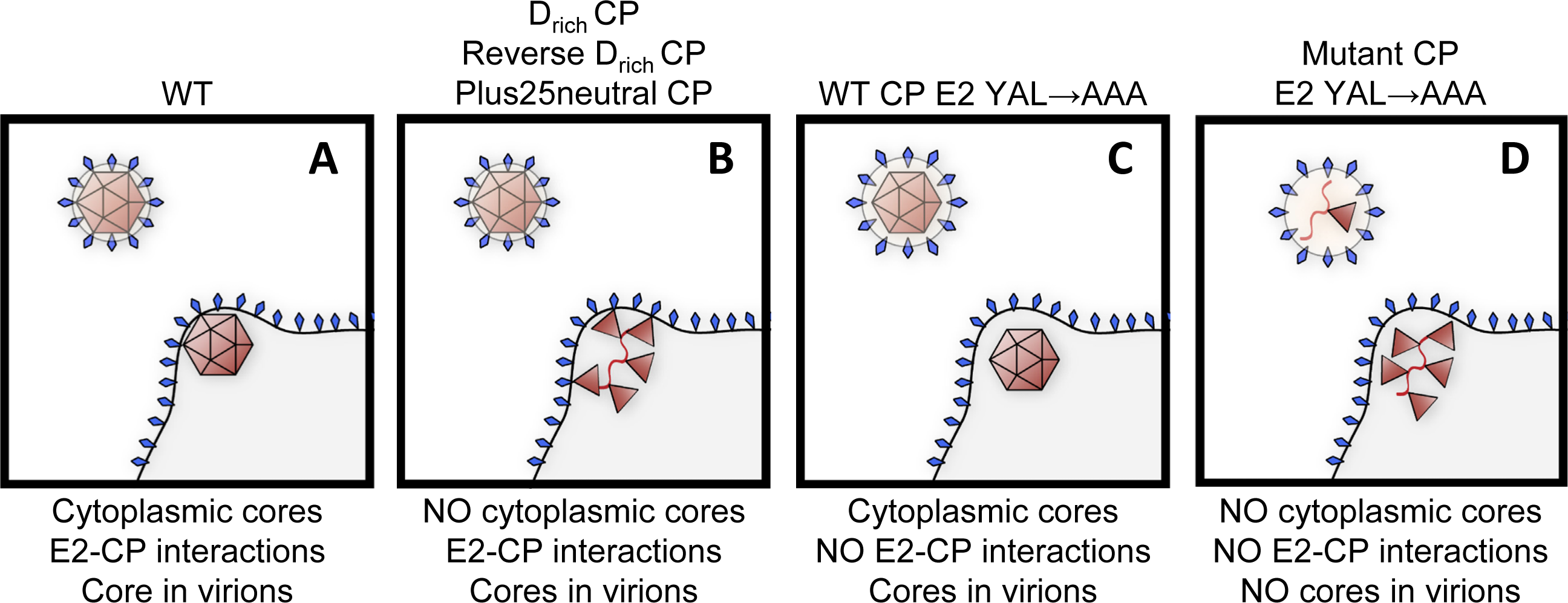
Schematic representation of the different ways cores or capsid protein (CP) interact with the cytoplasmic domain of E2 (cdE2) when the interacting proteins are mutated. (A) Nucleocapsid-directed assembly, the prevailing model of CPcdE2 interactions at the plasma membrane. The cytoplasmic core is thought to interact with cdE2 at the plasma membrane resulting in budding of a virus particle. (B) Glycoprotein-directed assembly. Core formation has been determined to be nonessential for virus particle release. In this case, CP and genomic RNA are still packaged in the released virions presumably by CP-RNA complexes interacting with cdE2 at the plasma membrane. (C) No nucleocapsid core-E2 interactions. cdE2 interactions with CP have been diminished by mutating the YxL motif in cdE2 resulting in a reduction in total particles and viral titer. (D) When both core formation and cdE2-CP interactions are diminished, we found that CP packaging decreases significantly.

### Cores can form in virus particles when cytoplasmic core formation is abrogated

As previously mentioned, other groups have described both N-terminal and Cterminal CP mutants that are unable to form stable cytoplasmic cores but often still release infectious virus particles (9, 12, 17–19, 26, 29–31, 33). The ability to isolate cores from virus particles was examined in some of these CP mutants and showed that the CP mutants were still unable to form cores (12, 19, 30, 31). When examining the CP mutants in this study, we found that even though the CP mutants were unable to form cytoplasmic cores, infectious virus was assembled and these particles had similar sedimentation profiles. When the glycoproteins were stripped from these particles, the CP migrated into the gradient for all of the mutants (Figure 5). This result is similar to what was demonstrated for the Sindbis (SINV) CP mutant TC-186; this mutant had cytoplasmic cores that were unstable to gradient centrifugation, but cores isolated from virus particles were stable (33). These authors suggest a maturation step. We go one step further and show cdE2 makes CP assembly competent since Plus25neutral can disassemble and reassemble into cores or core-like structures, similar to WT CP. D_rich_ and ReverseD_rich_ do not reassemble, most likely because the interaction with vRNA is weak due to the change in charge. It is interesting to note that the only examples of viruses that are able to form cores in virus particles in the absence of cytoplasmic core formation are insertion mutants-either at the N-terminus or in the C-terminal domain, although it is possible that some previously described mutants would have the same phenotype if cores from virions were examined. Interestingly, there has been an example of an alphavirus in the literature that has a contains a duplication in the CP N-terminus and still produces infectious virus, however, core formation has not been studied in this mutant (41).

### Packaging of CP into virions in the absence of cytoplasmic core formation is dependent on the interaction of CP with cdE2

When other labs have examined budding of alphaviruses via mutational analysis, the studies have focused on mutating either the CP or E2 where the proteins were proposed to interact or do interact (6, 8, 10, 13, 38). When there is a WT CP but a mutation in the E2 glycoprotein where it interacts with the core, the released virus particles still package cores but less efficiently than in the presence of WT E2 based on the reduction in viral titer (Figure 8C). And if there is a CP mutation and WT E2, infectious virus has been observed (Figure 8B). The CP is multi-functional in that it interacts with the viral genome with one domain leading to core formation, and interacts with E2 with the other domain. It is clear that the different functions of CP impact each other to different extents, but it is unclear what happens when both core formation and E2-CP interactions are disrupted concurrently. In this study, we hypothesized that the interaction of cdE2 with CP that is unable to form cytoplasmic cores rescues core formation in the virus particle either during or after budding. To test this hypothesis, cdE2 was mutated to decrease CP-cdE2 interactions in the context of the CP mutations. We proposed that cores would no longer be formed in the virus particles without CP-cdE2 interactions, but we found something more surprising (Figures 8D and 8E). None of the CP mutants were detectable in the released virions, indicating that in the absence of both core formation and CP-cdE2 interactions, CP packaging is severely diminished (Figure 7 and Figure 8E). Since infectious virions are still released from cells infected with these viruses, it is likely that genome is being packaged into a membrane containing glycoproteins without CP. This is not unprecedented since it has been demonstrated that virions, called infectious microvesicles (iMVs), can be released from cells infected with clones that do not have a CP (32).

### Core formation in virus particles is dependent on cdE2-CP interaction

Previous studies have shown that there are differences in the structure of the core at different stages of the viral lifecycle (42–44). When comparing the structures of cytoplasmic cores, core in virions, and cores isolated from virions in VEEV, cytoplasmic cores and cores isolated from virions were expanded in size compared to cores in virus particles. Additionally, there are differences in the capsomere organization in the structures (42–44). Together, the differences in the structures suggests that there is significant reorganization of capsomeres during or after budding and also during or after removal of the glycoproteins. It has been suggested that this reorganization is mediated either by the lipid bilayer or the viral glycoproteins (42, 43). When cytoplasmic cores are not formed, it has been proposed that CP packaging and core formation in virus particles is rescued by the CP-cdE2 interactions (19, 30, 31, 33). The results with the insertion mutants in this study and previously with the TC-186 mutant are consistent with this hypothesis since cores were found in virus particles that were not present in the cytoplasm (33). Through both the core disassembly and reassembly assay, and the cdE2 mutations in the CP mutant backgrounds we have shown that CP-cdE2 interactions are in fact necessary for not only CP packaging in the absence of cytoplasmic core formation but for virion core formation (Figures 6 and 7). This result adds to our understanding of what is necessary during the budding process, and has implications for our understanding of virion and core disassembly upon infection of a new cell.

### Implications of this work on core disassembly

Some groups have suggested that the rearrangement that occurs from cytoplasmic cores to virion cores during budding is the step that primes the core for disassembly (8, 9). The results in this study and with the TC-186 mutant actually show that cores isolated from virus particles can be more stable than cores in the cytoplasm contradicting this presumption (33). Instead, it appears likely that binding of CP to cdE2 actually results in more stable CP-CP interactions that are preserved even when the glycoproteins are removed (6, 20). So this leads to the question of what primes the core for disassembly upon infection in a new cell? If the core actually becomes more stable in the virus particle and doesn’t fall apart upon removal of glycoproteins, more steps must be involved for there to be a productive infection in a new cell. It has been shown that cores isolated from virions have holes at the base of the hexons, and that these cores are also more RNase sensitive (42, 45). Additionally, it has been demonstrated that cores in virus particles are often not complete closed shells and that this information is lost when icosahedral averaging is imposed (4). Experiments have shown that ribosomes are involved in disassembly of the viral core but the CP sequence that binds to the ribosome is internal to the core, meaning that the core must at least partially disassemble for ribosomes to have access to the binding site (20, 46, 47). Some groups have proposed that this partial disassembly occurs via exposure of the core to the low pH of the endosome leading to weakening of electrostatic interactions in the C-terminal CP domains (20).

However, it is possible that the holes in the hexons and/or the incomplete core structures either allows exposure of the CP N-terminus, specifically the ribosome binding sequence, to the exterior of the core, or allows the ribosome to access the interior of the core leading to disassembly. It has been shown in Hepatitis B virus that the interior CP domain can be exposed to the exterior of the core to bind cellular proteins (48). More studies will be needed to determine whether any of these possibilities occur during a viral infection and the relative contributions of each mechanism. Here we have shown that the steps leading to disassembly of the core likely occur after budding rather than during budding, and likely after entry into a new cell since stable cores can be isolated from budded virus particles. Further studying disassembly and how CP-cdE2 interactions are involved would be beneficial to developing inhibitors that could block alphavirus infections, especially if there is a conserved mechanism among all alphaviruses.

## MATERIALS AND METHODS

### Cloning of CP mutations

All of the capsid protein (CP) mutants were made in the RR64 plasmid (49). The D_rich_ and Reverse D_rich_ CP mutants were made using the QuikChange Lightning Site-Directed Mutagenesis kit (Agilent, Santa Clara, CA). For the D_rich_ mutant, the sequence for the negatively-charged addition was amplified from a previously made CP mutant (called 10D) using the following primers: (F) 5’CTTCATCTAATACAGCTCACAACAG- TAAACATGGAAAACCAGGACGATGACCC-3’ and (R) 5’- CAATTGTGCCTTTAAC-

GTGAGCCG-3’ resulting in a 25 amino acid addition between AA1 and AA2 in the wild-type (WT) CP sequence. The insert sequence for the Reverse D_rich_ CP mutant was amplified from the WT CP sequence and inserted between AA1 and AA2 in the 10D CP sequence using the following primers: (F) 5’- AGCTCACAACAGTAAACATGGAAAAC- CAGAAGAAAAAGCC-3’ and (R) 5’- GTCTGGGTTGG-TATGTAATTTTTCTTCTTCTTAGCCTGTG-3’. The Plus25neutral mutant was generated via Gibson assembly using CloneAmp^TM^ 2X HiFi PCR mix (Takara Bio USA, Mountain View, CA) for backbone amplification, and NEBuilder^®^ HiFi DNA Assembly Master Mix (NEB, Ipswich, MA) for the Gibson assembly. The backbone sequence was amplified from the WT plasmid using the following primers: (F) 5’CACAGGCTGCCAAC- GCCAATAATTACATACCAACCCAGAC-3’ and (R) 5’- GGGGCATTGGCCTGGTTT-GCCATGTTTACTGTTGTGAGCT-3’. The insert sequence was ordered from GeneArt in a pMA-T vector (ThermoFisher Scientific, Waltham, MA), and amplified using Taq DNA polymerase with the following primers: (F) 5’- ACAACAGTAAACATGGCAAAC- CAGGCCAATGCCCCGACGCAAGCCAACGCCCAGCAGCAG-3’ and (R) 5’- GTCTGGGTTGGTATGTAATTATTGGCGTTGGCAGCCTGTG-3’. The sequence was inserted between AA1 and AA2 in the WT CP sequence. All cloning was confirmed by DNA sequencing of the mutated regions.

### Cloning of E2 mutations

Mutation of E2 residues 399YAL401 to AAA was achieved using the QuikChange Lightning Site-Directed Mutagenesis kit (Agilent, Santa Clara, CA). The following primers were used: (F) 5’- GAAAGTGCCTAACACCAGCCGCCGCGACGCCAGGAGCGGTGG- 3’ and (R) 5’- CCACCGCTCCTGGCGTCGCGGCGGCTGGTGTTAGGCACTTTC-3’.

Successful cloning was confirmed via DNA sequencing of the E2 protein sequence.

### Tissue Culture Reagents

All experiments were conducted in baby hamster kidney-21 cells, herein referred to as BHK cells. Cells were passaged in 1X minimal essential media (MEM) supplemented with 1% L-glutamine, 1% MEM non-essential amino acids (NEAA), 1% penicillin-streptomycin and 5% fetal bovine serum (FBS, Corning, Corning, NY). Virus prepared for purification by ultracentrifugation was grown in serum-free media (VP-SFM; Gibco, ThermoFisher Scientific, Waltham, MA) supplemented with 1% (v/v) L-glutamine, 1% (v/v) MEM NEAA, and 1% (v/v) penicillin-streptomycin or antibiotic-antimycotic (Corning, Corning, NY). Cells were incubated at 37°C and 5% CO_2_.

### Virus Preparation and Titering

Infectious virus was prepared from a cDNA clone. Each viral cDNA clone was linearized with Sac1 (NEB, Ipswich, MA), and transcribed into viral RNA *in vitro* using SP6 polymerase, and cap analog (NEB, Ipswich, MA). The viral RNA was electroporated (1500V, 25 μF, 200Ω) into BHK cells resuspended in phosphate-buffered saline (PBS, Corning, Corning, NY) in a 2mm gap-width cuvette. Cells were plated in 1X MEM supplemented with NEAA, penicillin-streptomycin, L-glutamine, and 5% FBS. Media was harvested at 48 hours post electroporation (P0 virus), and cellular debris removed by centrifuging at ∼1100xg and 4°C for 5 minutes. P1 virus stocks were generated by infecting a confluent monolayer of BHK cells with 1mL of the respective P0 virus. Media was harvested at 24hpi and the cellular debris removed by centrifuging at ∼1100xg and 4°C for 5 minutes.

Viral titers were determined via standard plaque assay on BHK cells. Briefly, serial dilutions of viral stocks or sucrose fractions were added to 70-90% confluent BHK monolayers in 6or 12-well plates, and adsorbed at room temperature for 1 hour with rocking.

Cells were overlayed with 2% low-melt agarose mixed 1:1 with complete 2X MEM containing 10% FBS and then fixed with formaldehyde at 48 hours post infection and plaques visualized via crystal violet staining. Viral titers are presented as plaque forming units (PFUs) per mL.

Infections were conducted at the multiplicity of infection (MOI) indicated in each figure legend. Virus was adsorbed to the BHK cells at the indicated MOI for 1 hour at room temperature with rocking. The inoculum was removed, the cells washed with PBS, and fresh media added to the cells before incubating them at 37°C and 5% CO_2_ until harvest. In Figures 2A and 3A, P0 virus stocks were used to infect cells at an MOI=1 for 24 hours. In figure 2B, cells were infected with P1 virus stocks at an MOI=0.01 for 20 hours or 24 hours, or an MOI=1 for 24 hours (MOI and infection time were consistent within an experiment). For Figures 7A and 7B, cells were infected with P1 virus stocks at an MOI=0.0001 for 48 hours. To prepare cell lysates, media was removed from the cells and the cells were lysed in BHK cell lysate buffer (10mM Tris pH 7.4, 20mM NaCl, 0.4% deoxycholic acid, 1% NP-40, and 1mM EDTA).

### SDS-PAGE and Western Blotting

All SDS-PAGE gels are 12% bis-acrylamide gels containing trichloroethanol (TCE, Santa Cruz Biotechnology, Dallas, TX) at 0.5% v/v. Western blots were probed with α- CP or α-RRV E2 primary antibodies at a 1:5000 dilution in 2% milk in TBST followed by AlexaFluor^TM^750 goat α-rabbit secondary antibody (Invitrogen A21039, Carlsbad, CA) at a 1:20,000 dilution in 2% milk in TBST. All images were collected on a ChemiDoc system (BioRad, Hercules, CA) and analyzed using Fiji (50).

### Transmission Electron Microscopy (TEM)

Five microliters of purified virus or pelleted cores were placed on a Formvar and carboncoated 300 mesh grid (Ted Pella Inc., Redding, CA) and stained with 1% uranyl acetate. If samples were very concentrated, they were diluted 1:10 in HN buffer (20mM HEPES, 150mM NaCl, pH 7.4) prior to staining. Stained grids were viewed using a JEOL 1010 transmission electron microscope (Tokyo, Japan) operating at 80kV, and images captured at 25000X using a Gatan Megascan 794 charge-coupled-device camera (Gatan Inc., Pleasanton, CA). Images were converted into TIFF format using Fiji (50).

### Purification of Virus

Confluent BHK cells were infected at an MOI=0.1 with absorption for 1 hour at room temperature with rocking. After absorption, the inoculum was removed, the cells washed three times with PBS, and then incubated in VP-SFM supplemented with L-glutamine, NEAA, and penicillin-streptomycin or antibiotic-antimycotic for 32 hours at 37°C and 5% CO_2_. The viral supernatant was clarified at ∼1100xg for 5 minutes at 4°C. Clarified supernatant was loaded over 3mL of a 27% w/v sucrose (in HN buffer) cushion and pelleted at ∼186,000xg for 2 hours at 4°C. The supernatant was poured off and each pellet resuspended in 200µL HN buffer or TNE (10mM TrisCl, 10mM NaCl, 20mM EDTA, pH 7.5) buffer. When the virus particles were further purified over a sucrose gradient, 20- 50% w/v sucrose gradients were generated on the gradient master 107 system (Bio- Comp, New Brunswick, Canada) using the setting Short Sucr 20-50% wv 1ST. The two pellets for each virus were combined and diluted to 1mL in HN buffer. 800µL was removed from the top of each gradient before loading 1mL of virus to the top of each gradient. The gradients were centrifuged at ∼247,000xg and 4°C in an SW41 rotor for 3 hours. Following centrifugation, 1mL fractions were collected from the top of each gradient and saved for further analysis.

### Isolating Cores from Cells

A protocol was developed based on Lopez *et al*. (38). Infected one 100mM dish of BHK cells per virus at an MOI=0.1 or MOI=1 for 20-24 hours. At 20-24hpi, the media was removed and discarded from the cells, and the cells washed with 1X PBS. The cells were manually scrapped from the dish into the PBS, transferred to a sterile tube, the cells pelleted at ∼1100xg and 4°C for 5 minutes, and the supernatant discarded. The cells were resuspended by pipetting in fresh PBS, pelleted at ∼1100xg and 4°C for 5 minutes, and the supernatant discarded twice more. Following the final PBS wash, the cells for each infection were resuspended in 1mL TNE buffer with pipetting, and then incubated on ice for 20 minutes. After the incubation, 200µL of 20% TritonX-100 was added to each tube of resuspended cells, and gently vortexed for ∼10 seconds to lyse the cells. The tubes were centrifuged at 1500xg and 4°C for 10 minutes to pellet the nuclei. The supernatant for each sample was transferred to a new tube and stored on ice until the sample was loaded on the gradient. 10-40% w/v sucrose (in TNE-T buffer) gradients were prepared on the gradient master 107 system (BioComp, New Brunswick, Canada) using the setting Short Sucr 10-40% wv 1ST. 800µL was removed from the top of each gradient, prior to loading 1mL of the supernatant saved on ice to the top of each gradient. The gradients were centrifuged at ∼175,000xg and 4°C in an SW41 rotor for 2.5 hours.

Following centrifugation, 1mL fractions were collected from the top of each gradient and saved for further analysis.

### Isolating Cores from Virions

Cores were isolated similarly to (19). Briefly, virus particles were purified by pelleting through a sucrose cushion as in the section “Purification of Virus” with the pellets resuspended in TNE buffer. The two pellets for each virus were combined and then diluted 1:2 in water. NP-40 was added to each diluted virus sample to a final concentration of 1% and mixed by pipetting up and down, followed by a 10 minute incubation on ice. Linear 10-40% w/v sucrose gradients were prepared on the gradient master 107 system (BioComp, New Brunswick, Canada) using the setting Short Sucr 10-40% wv 1ST.

600µL was removed from the top of each gradient prior to loading 800µL of lysed virus particles to the top of each gradient. The gradients were centrifuged at ∼175,000xg and 4°C in an SW41 rotor for 2.5 hours. Following centrifugation, 1mL fractions were collected from the top of each gradient and saved for further analysis.

### Purifying Cores through a Sucrose Cushion

Virus particles were purified by pelleting through a sucrose cushion as in the section “Purification of Virus” with the pellets resuspended in TNE buffer. Pelleted virus from three 150mm dishes (∼1.26x10^8^ BHK cells total) was resuspended in 400µL TNE buffer and diluted 1:2 in water. 80µL of 10% NP-40 was added, the sample mixed by pipetting up-and-down, and the samples incubated on ice for 10 minutes. 800µL of each sample was loaded on top of a 20% w/v sucrose cushion in HN buffer and centrifuged at 110,000xg and 4°C for 2 hours in a SW55 rotor. Following centrifugation, 1mL was removed from the top of the sucrose cushion for the top of cushion sample, the rest of the sucrose cushion saved, and then the pellets resuspended in 50µL HN buffer.

### Disassembly and Reassembly Assay

The concentration of cores was determined by nanodrop with the assumption that 1Abs=1mg/mL. 15µg of cores was diluted into 30µL using HN buffer, and then measured on the Zetasizer as described below in the DLS section. 5.57µL of the sample was removed and 1.90µL of 5M NaCl was added to the remaining sample to bring the final salt concentration up to 500mM. The 500mM sample was measured on the Zetasizer as described below in the DLS section. 6.00µL of the sample was removed and 47.44µL of HN buffer was added to the remaining sample to dilute the salt concentration back down to 150mM by pipetting up and down. The diluted sample was measured on the Zetasizer as described below in the DLS section and then 20µL of the sample saved.

The saved samples were for the agarose gel-shift analysis.

### Dynamic Light Scattering (DLS)

DLS measurements were conducted at 25°C in a Malvern Zetasizer Nano ZS instrument with the attenuator set to 9 and the measurement window set to 4.2mm (Malvern Panalytical, Malvern, UK). A Zen2112 quartz cuvette (Malvern Panalytical, Malvern, UK) with a 3mm path length was used for the experiments. All samples were measured three times with each measurement consisting of 10 runs with a run averaged over 10 seconds. The data is graphed with average hydrodynamic diameter (nm) on the x-axis and normalized intensity on the y-axis. Normalized intensity was calculated using the intensity percentage for the data points, the derived count rate, and accounting for the dilution factor. The DLS data was analyzed and visualized using Microsoft Excel 2013 and GraphPad Prism.

### Agarose gel-shift assay

For the 150mM salt samples, 5.57µL of sample was combined with 1.2µL 6X DNA loading dye (NEB, Ipswich, MA); for the 500mM salt samples, 6µL of sample was combined with 1.2µL of 6X DNA loading dye; for the 150mM salt samples, 20µL of sample was combined with 4µL 6X DNA loading dye. Volume of samples were varied according to the dilution factor. The samples were loaded on a 0.8% agarose gel in TBE buffer (45mM Tris-Base, 0.55% Boric Acid, 1mM EDTA), the gel run at 100V (constant) for 30 minutes, and then nucleic acid detected using ethidium bromide. The gel was then soaked in gel drying solution (20% methanol, 10% acetic acid) for 15 minutes, dried for 1 hour at 70°C on a BioRad Model 583 gel dryer (BioRad, Hercules, CA), and protein visualized with Coomassie staining. Images were captured on the BioRad ChemiDoc MP Imaging System (BioRad, Hercules, CA) and the images were processed in Fiji (50).

## ACKNOWLEDGMENTS

This work was funded by a National Institutes of Health training grant T32GM109825 (JMB) and a Milton Taylor Virology Fellowship (JMB). The content is solely the responsibility of the authors and does not necessarily represent the official views of the National Institutes of Health.

We would also like to thank the IU Nanoscale Characterization Facility for access to instrumentation. We would like to acknowledge A. Moore and the IU Virology group for productive scientific discussions and suggestions related to this work.

